# O-Pair Search with MetaMorpheus for O-glycopeptide Characterization

**DOI:** 10.1101/2020.05.18.102327

**Authors:** Lei Lu, Nicholas M. Riley, Michael R. Shortreed, Carolyn R. Bertozzi, Lloyd M. Smith

**Affiliations:** Department of Chemistry, University of Wisconsin, Madison, WI 53706; Department of Chemistry, University of Stanford, Stanford CA 94305; Howard Hughes Medical Institute, Stanford CA 94305

## Abstract

We report O-Pair Search, a new approach to identify O-glycopeptides and localize O-glycosites. Using paired collision- and electron-based dissociation spectra, O-Pair Search identifies O-glycopeptides using an ion-indexed open modification search and localizes O-glycosites using graph theory and probability-based localization. O-Pair Search reduces search times more than 2,000-fold compared to current O-glycopeptide processing software, while defining O-glycosite localization confidence levels and generating more O-glycopeptide identifications. O-Pair Search is freely available: https://github.com/smith-chem-wisc/MetaMorpheus.

---

Mass spectrometry (MS) is the gold standard for interrogating the glycoproteome, enabling the localization of glycans to specific glycosites.^1–3^ Recent applications of electron-driven dissociation methods have shown promise in localizing modified O-glycosites even in multiply glycosylated peptides^4^. Yet, standard approaches for interpreting tandem MS spectra are ill-suited for the heterogeneity of O-glycopeptides. Perhaps the most challenging problem for O-glycopeptide analysis is mucin-type O-glycosylation, which is abundant on many extracellular and secreted proteins and is a crucial mediator of immune function, microbiome interaction, and biophysical forces imposed on cells, among others^5^. Mucin-type O-glycans are linked to serine and threonine residues through an initiating N-acetylgalactosamine (GalNAc) sugar, which can be further elaborated into four major core structures (cores 1-4) or remain truncated as terminal GalNAc (Tn) and sialyl-Tn antigens^6^. These O-glycosites occur most frequently in long serine/threonine rich sequences (**Supplementary Fig. 1**), such as PTS mucin tandem repeat domains, which exist with microheterogeneity defined by a large number of potential O-glycans^7^. The number of serine and threonine residues present in glycopeptides derived from mucin-type O-glycoproteins, combined with the consideration of dozens of potential O-glycans at each site, leads to a combinatorial explosion when generating databases of theoretical O-glycopeptides to consider for each tandem MS/MS spectrum (**Supplementary Note 1**).

Current O-glycoproteomic analysis pipelines are unable to search for multiply O-glycosylated peptides within reasonable time frames even for simple mixtures of O-glycoproteins, much less for proteome-scale experiments. Recent efforts to combat this search time issue have forgone site localization for the more expedient option of identifying only the total glycan mass on a peptide backbone^8^. While effective at lowering time costs, this sacrifices valuable information about site-specific modifications – which is often the goal of intact glycopeptide analysis in the first place. Open modification searches and combinations of peptide database searching with *de novo* glycan sequencing have also recently been reported, but neither address the time issues that challenge analysis of highly modified O-glycopeptides.^9,10^ Moreover, electron-driven dissociation methods are required to localize O-glycosites^11^, yet current software tools fail to capitalize on combinations of collision-based and electron-based fragmentation spectra that are acquired for the same precursor ion. This is coupled with a general lack of ability to confidently localize glycosites within multiply glycosylated O-glycopeptides.

Here, we describe the O-Pair Search strategy implemented in the MetaMorpheus platform^12^ to provide a pipeline for rapid identification of O-glycopeptides and subsequent localization of O-glycosites using paired collision- and electron-based dissociation spectra collected for the same precursor ion (**Fig. 1a**). O-Pair Search first uses an ion-indexed open search^13^ of higher energy collisional dissociation (HCD) spectra to rapidly identify combinations of peptide sequences and total O-glycan masses, which are generated through combinations of entries in an O-glycan database. Graph-theoretical localization^14–17^ then defines site-specific O-glycan localizations using ions present in EThcD spectra (electron transfer dissociation with HCD supplemental activation) (**Fig. 1b**). Peptide backbone fragments (b/y-type ions) rarely retain glycan mass during HCD fragmentation, making them good candidates for an ion-indexed search, while retention of intact glycans on c/z^•^-fragments in EThcD spectra enable confident localization,^11^ as exemplified in the paired HCD-EThcD spectra for the quadruply glycosylated peptide in **Fig. 1c**. Localization is followed by localization probability calculations using an extension of the phosphoRS^18^ algorithm used for phosphosite localization (max score of 1), in addition to scoring of fine scoring (which includes calculation of Y-type ions) and false discovery rate calculations performed separately for O-glycosylated and non-modified peptides.

**Figure 1.**
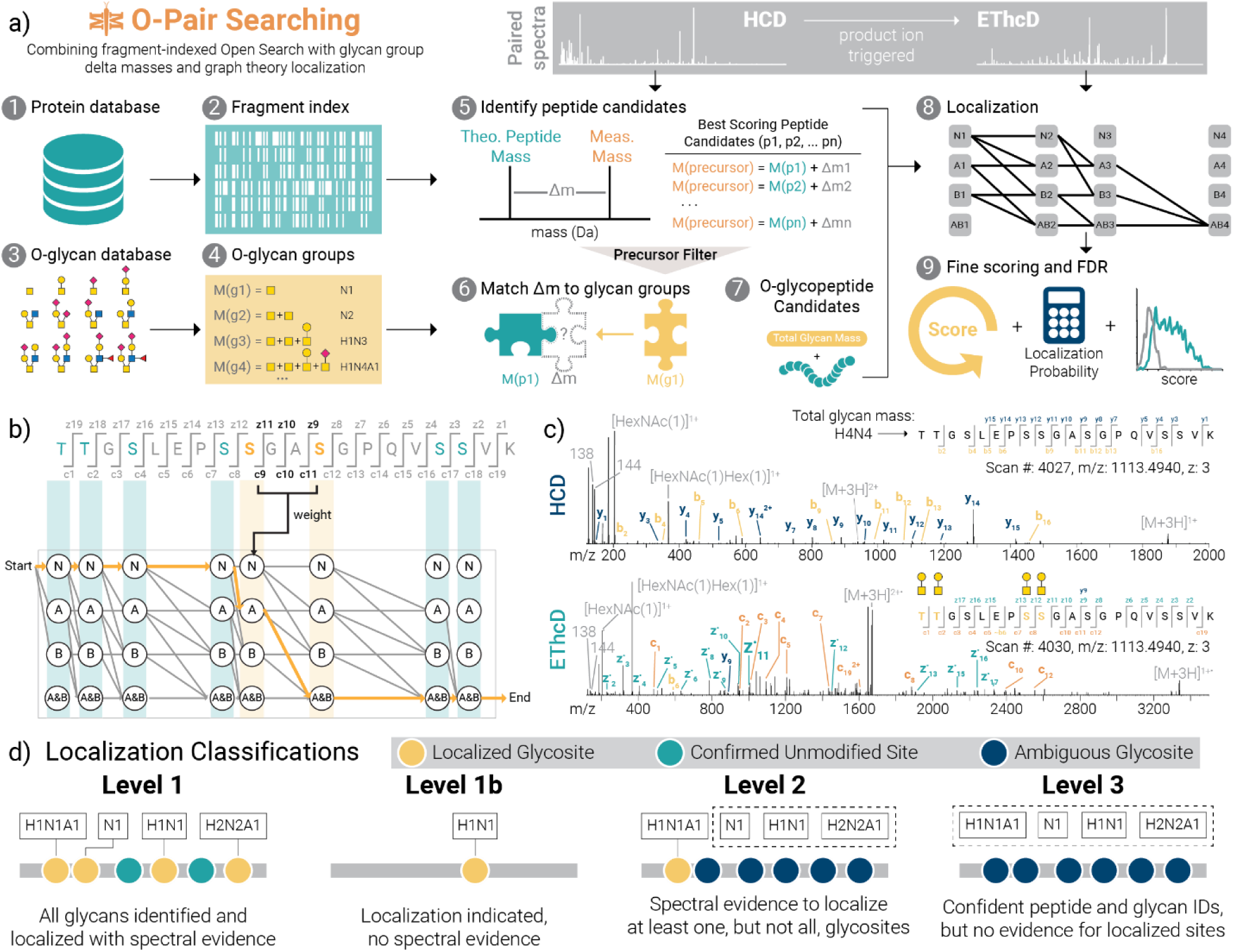
O-Pair Search through MetaMorpheus for fast and confident identification of O-glycopeptides. **a)** The workflow describes processing steps in the O-Pair Search strategy, which generates a fragment ion index [1, 2] and O-glycan groups [3, 4] from user defined protein and O-glycan databases, respectively. Using an ultrafast, fragment-index-enabled open modification search [5] paired with a match of delta masses to aggregate glycan mass combinations [6] enables identification of O-glycopeptide candidates from HCD spectra [7]. Paired EThcD spectra are then used for graph theory-based localization calculations to rapidly assign modification sites for all glycans comprising the O-glycan group [8]. Finally, more detailed re-scoring of spectra, localization probability calculations, and false discovery rate corrections are performed before returning identifications to the user [9]. **b)** A demonstration of graph theory-based localization using a hypothetical example of an O-glycopeptide TTGSLEPSSGASGPQVSSVK from human mucin-type O-glycoprotein CD43 (leukosialin), which has 8 possible O-glycosites. Here we consider how graph theory determines O-glycosites using c/zdot fragments present in EThcD spectra when two glycans (termed A and B for the sake of demonstration) are presented as modifications. **c)** An example of paired HCD and EThcD spectra for quadruply-O-glycosylated TTGSLEPSSGASGPQVSSVK, showing a Level 1 identification where all calculated glycan mass shifts can be confidently localized to discrete residues. Note, no fragments in the HCD spectrum retain any glycan masses. Rather, the thorough peptide backbone fragmentation without glycan retention shows how the sequence was confidently retrieved with a defined mass shift matching a combination of O-glycans. The subsequent EThcD spectrum then enables localization of all 4 O-glycosites (gold) even with the presence of 4 other unmodified potential sites. D) O-Pair Search defines levels of localization for each GlycoPSM. A Level 1 assignment indicates that all glycans can be unambiguously localized to single S or T residues using spectral evidence, while Level 1b also indicates localization in instances when spectral evidence is lacking (e.g., only one possible modification site). Level 2 localizations have at least one glycan, but not all, localized to a single S or T. Level 3 GlycoPSMs include the remaining pool of identifications, where peptide sequence and glycan aggregate mass are confidently assigned, but no individual glycan can be localized to a specific residue. Note, “H”, “N”, and “A” represent hexose, HexNAc, and Neu5Ac, respectively.

We also introduce here the concept of Localization Levels, which is the culmination of the O-Pair Search (**Fig. 1d**). Inspired by early adoption of class levels for phosphopeptide localizations^19^ and more recently for proteoforms^20^, we developed this classification system to more accurately describe the quality and confidence of glycopeptide and glycosite identifications. *Level 1* glycopeptide identifications indicate that all glycans identified in the total glycan mass modification are localized to specific serine and threonine residues with a localization probability > 0.75. Glycopeptides with glycosite assignments with localization probabilities < 0.75 are assigned as *Level 1b*, even though they are still identified as localized by the graph theory approach. *Level 1b* assignments also occur when a glycosite is assigned without the presence of sufficient spectral evidence (e.g., fragments cannot explain a glycosite, but the sequence contains only one serine or threonine). We currently borrow the 0.75 cutoff from phosphopeptide precedents^19^; empirical determination of localization cutoffs will likely need to be determined in future work using libraries of synthetic glycopeptide standards, as has been done with phosphopeptides^21^. That said, such libraries are currently difficult to generate. *Level 2* assignments occur when at least one glycosite is assigned a glycan based on spectral evidence, but not all glycans can be assigned unambiguously. *Level 3* identifications represent a confident match of glycopeptide and total glycan mass, but no glycosites can be assigned unambiguously. Indeed, *Level 3* glycopeptides (such as those reported in HCD-only methods by default^8^) are still useful to note the presence of glycosylated residues somewhere in a given sequence. Our classification system provides a straightforward approach to qualify glycoproteomic datasets without having to exclude confident identifications that have no site-specific information. In addition to Localization Level assignments, O-Pair Search also reports the ratio of oxonium ions (known to help distinguish glycan type^22^), the presence of N-glycosylation sequons to identify potentially confounding assignments, matched peptide and glycan fragment ion series and their intensities for each of the paired spectra, and localization probabilities for all sites, both localized and not.

We first compared O-Pair Search to Byonic, the most commonly used O-glycopeptide identification software^23^. Byonic, which uses a look-up peaks approach to speed up search times relative to traditional database searching^24^, can also search HCD and EThcD spectra, although it is agnostic of paired spectra originating from the same precursor. To benchmark performance, we used a recently published dataset^11^ of O-glycopeptides from mucin glycoproteins using a combination of trypsin and the mucin-specific protease StcE, which cleaves only in glycosylated mucin domains^25^. This data originates from sequential digestion of four recombinant mucin standards (CD43, MUC16, PSGL-1, and Gp1ba), using StcE to cleave mucin domains followed by N-glycan removal with PNGaseF and tryptic digestion. We initially searched a file with HCD and EThcD paired spectra from this dataset, and a glycan database of 12 common O-glycans was used for these searches^8^. O-Pair Search identified more localized (**Fig. 2a**) and total (**Fig. 2b**) O-glycopeptide spectral matches (GlycoPSMs) than Byonic when allowing either 2 or 3 glycans per peptide (**Supplementary Fig. 2** and **Fig. 2**, respectively). This holds true even when relaxing the scoring thresholds used to obtain confident Byonic identifications (**Supplementary Fig. 3**). Note, all O-Pair Search identifications represent two spectra from an HCD-EThcD spectral pair. Conversely, Byonic is agnostic to paired scans, meaning identifications can come from HCD and EThcD spectra that were collected for the same precursor (pair) or from spectra identified separately from their paired counterpart.

**Figure 2.**
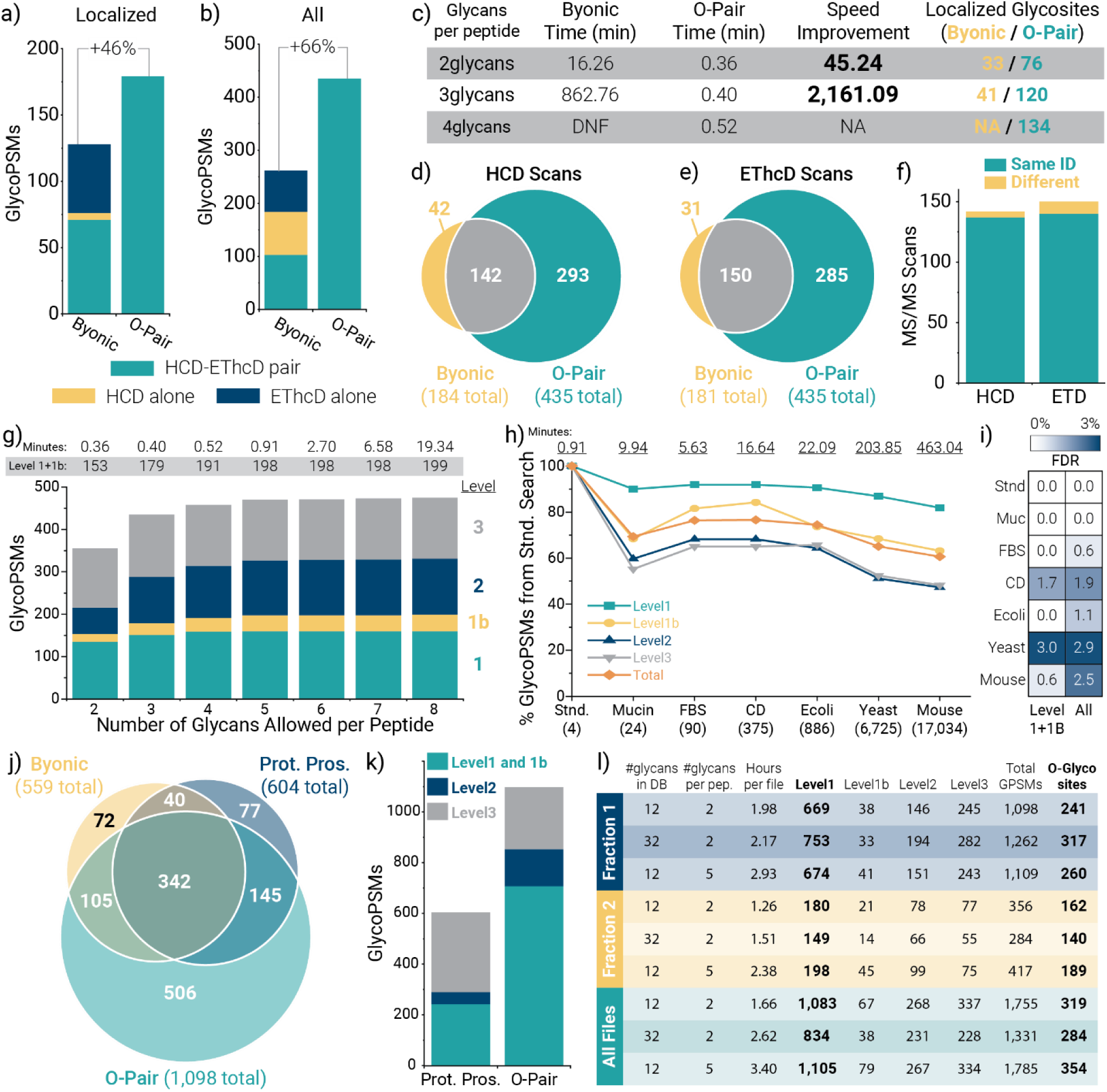
Performance of O-Pair Search for O-glycopeptide characterization. Comparing the number of **a)** localized and **b)** total glycopeptide spectral matches (GlycoPSMs) returned from Byonic and from O-Pair Search for HCD-pd-EThcD data collected from StcE digestions of four recombinant mucin standards. Note, only Level 1 and 1b identifications are considered for the localized O-Pair Search data. Byonic identifications are grouped into HCD-EThcD pairs (where paired scans identified the same O-glycopeptide), HCD alone, and EThcD alone. The latter two cases are where an identification came only from an HCD scan or EThcD scan, but the other spectrum in the pair did not return a hit. O-Pair Search improves the number of localized and total identifications by 46% and 66% over Byonic, respectively. **c)** The table compares the search times required for Byonic and O-Pair Search when considering 2, 3, and 4 glycans per peptide. Note, the 4 glycans per peptide for Byonic was canceled after approximately 33,000 minutes of search time (∼3.5 weeks) because it had not advanced in reported search progress for over one week. The number of localized glycosites identified by the searches is also provided for comparison. In addition to more than doubling the number of total identified spectra, O-Pair Search identified the majority of scans that Byonic returned as GlycoPSMs for both **d)** HCD and **e)** EThcD scans, and **f)** the overwhelming majority (∼95%) of the shared identified scans mapped to the same glycopeptide. **g)** O-Pair Search enabled consideration of more glycans per peptide while keeping search times reasonable. **h)** O-Pair Search also allowed the use of several different protein database backgrounds much larger in size without untenable search time increases. **i)** Use of entrapment databases with proteins not present in the sample did not inflate false discovery rates above approximately 1-3%. **j)** O-Pair Search was used to process files from a published urinary O-glycopeptide study that previously reported Protein Prospector (Prot. Pros.) and Byonic results. O-Pair Search nearly doubled the total number of GlycoPSMs from either search engine, identifying ∼90% of spectra shared by the two search algorithms while providing an additional 506 GlycoPSMs not reported by either. **k)** Protein Prospector reports localized glycosites, which we converted into our Localization Level system and compared with O-Pair results. **l)** Results from several O-Pair searches of Fraction 1 (three files), Fraction 2 (two files), and all ten files available from the urinary O-glycopeptide study.

Importantly, O-Pair Search dramatically decreased search times, with ∼45-fold and ∼2,100-fold faster searches than Byonic when considering 2 or 3 glycans per peptide, respectively (**Fig. 2c**). O-Pair Search required approximately 30 seconds to complete a search considering 4 glycans per peptide, while the Byonic search was terminated after the search failed to complete in over 33,000 minutes (∼3.5 weeks). Improvements in search speed are accompanied by ∼2-3-fold increases in the number of localized glycosites identified. In addition to more than doubling the number of total identified spectra, O-Pair Search identified the majority of spectra that Byonic returned as GlycoPSMs for both HCD (**Fig. 2d**) and EThcD (**Fig. 2e**) scans, and the overwhelming majority (∼95%) of the shared identified scans mapped to the same glycopeptide (**Fig. 2f**). These searches were completed using a FASTA file containing sequences only for the four mucin standards, which highlights the impracticality of O-glycopeptide searches in Byonic for complex mixtures. Moreover, O-Pair Search performed localization calculations and reported Localization Levels within the reported search time while Byonic spectra had to be further processed after the search to obtain localization information.

The ability to rapidly search O-glycopeptide data allowed us to vary the number of O-glycans to consider per peptide for easy evaluation of optimal search conditions. **Fig. 2g** shows that search times remain less than a minute when considering 5 glycans per peptide, while up to 8 glycans can be considered per peptide in searches requiring less than 20 minutes. Allowing for more glycans per peptide does not change the spectral assignments to various glycopeptides (**Supplementary Fig. 4**), indicating the robustness of O-Pair Search identifications. The number of non-modified identifications remained similarly constant (**Supplementary Fig. 5**).

Evaluating retention time rules further supports O-Pair Search identifications, where glycopeptide identifications containing 0, 1, and 2 sialic acids on the same peptide backbone have predictable elution time shifts (**Supplementary Fig. 6**)^26,27^. O-Pair Search localization of different glycosites also enabled visualization of chromatographically resolved glycopeptide positional isomers (**Supplementary Fig. 7**). Interestingly, processing of this published dataset to evaluate the best fragmentation conditions for O-glycopeptides generated the same overall conclusions as the previously reported Byonic searches, although the differences between different supplemental activation energies for EThcD appear more subtle than before (**Supplementary Fig. 8**). Note, these searches were completed using 16 cores, but similar performance can also be achieved on most standard computing systems using fewer cores (**Supplementary Fig. 9**). Overall, this method enabled characterization of dozens of glycosites on each glycoprotein in the mixture (**Supplementary Fig. 10**).

We also evaluated O-Pair Search search times and false discovery rates using several entrapment protein databases with varying complexity (**Fig. 2h**). A description of the databases used for benchmarking is provided in Supplemental Note 2; briefly, databases were designed to represent different proteome backgrounds not present in the sample (true negatives), with the four mucin standard target sequences (true positives) appended. Entrapment backgrounds ranged from 20 canonical human mucins to the entire mouse proteome. Search times for the mucin, FBS, cell surface glycoprotein, and E. coli entrapment databases (all with < 1,000 entries) remained under ∼20 minutes when using 16 cores, while the yeast and mouse entrapment proteomes took ∼3.4 and ∼7.7 hours. Still, this is approximately half the time Byonic required for a far less complex search (**Fig. 2c**). Sensitivity, as measured by the number of O-glycopeptide identifications, varied with the entrapment backgrounds, which was also evident for non-modified peptide identifications (**Supplementary Fig. 5**). This highlights the known issue of proper database size selection in glycoproteomics^28^, which can be more thoroughly explored for O-glycoproteomics now that O-Pair Search enables reasonable search times. Importantly, Level 1 peptides were the least affected, supporting their high confidence assignments. O-Pair Search maintained acceptable false discovery rates (0-3%) even when challenged with these entrapment databases (**Fig. 2i**), performing well compared to previous reports^29,30^.

Finally, we applied O-Pair Search to a large dataset of urinary O-glycopeptides, which has been analyzed in a number of studies^31–34^. The raw data for this dataset represents glycopeptides purified from urine from three healthy male donors using affinity chromatography with wheat germ agglutinin and is available through the MassIVE repository (MSV000083070). Pap et al.^31^ provide identifications from Protein Prospector and Byonic for EThcD scans from Fraction 1 (the “shoulder fraction”, three raw data files available) and Fraction 2 (the “GlcNAc fraction”, two raw data files available). We searched Fraction 1 with O-Pair Search using the entire human proteome database (∼20,300 entries) with 2 glycans considered per peptide from the 12 common O-glycan database used above, and we compared the results to the reported identifications for the other two search engines (**Fig. 2j**). Because this dataset had the potential to harbor N-glycopeptides as well, we filtered out all identifications that included an N-sequon from our O-Pair Search results. Even so, O-Pair Search nearly doubled the total number of GlycoPSMs from either search engine. Of the 382 spectra identified by both Protein Prospector and Byonic, O-Pair Search identified ∼90% of them (342 spectra) while providing an additional 506 GlycoPSMs not reported by either. Of the total 1,287 spectra identified as GlycoPSMs, O-Pair Search identified ∼85% of them (1,098 spectra). The original study reported a predominance of sialylated glycopeptides, which is recapitulated by O-Pair Search with >97.5% of GlycoPSMs (1,071 of 1,098) containing a sialic acid. When comparing identifications from the 342 scans identified in all three search algorithms, all return the same glycopeptide sequence. Protein Prospector reports a Site Localization In Peptide (SLIP) score^35^ for modification sites that we used to convert identifications to our Localization Level scheme (**Fig. 2k**). O-Pair Search reports more Level 1 and 1b O-glycopeptide identifications than the total number of Protein Prospector GlycoPSMs, and the proportion of localized and partially localized identifications (Levels 1-2) is more favorable with O-Pair Search. Similar trends hold for Fraction 2 (**Supplementary Fig. 11**).

We expanded our analysis of this dataset to explore the use of a larger glycan database (32 glycans vs 12) and the effect of searching with more glycans allowed per peptide (5 vs 2). **Fig. 2l** compares results from these different search parameters for Fraction 1, Fraction 2, and all 10 files available for download from the urinary O-glycoproteome dataset. In Fraction 1, The larger O-glycan database boosted identifications for Fraction 1, but lowered identifications in Fraction 2 and the entire dataset as a whole. This indicates that Fraction 1 likely harbored glycopeptides with more diverse glycans while the majority of the dataset did not. Conversely, considering more glycans per peptide provided slight benefits in all cases. By requiring only a few hours to perform a whole proteome-search with a variety of glycopeptide possibilities, O-Pair Search provides a flexible platform to explore O-glycoproteomics data. A recently published large dataset of human urinary O-glycopeptides identified ∼1,300 intact O-glycopeptides but was not able to report localized O-glycosites because of the reliance on HCD^36^. They did use EThcD to report 127 O-glycopeptides following their HCD study to confirm identifications, but we were unable to find their raw data publicly available to search. When considering only Level 1 and 1b GlycoPSMs, our results represent 447 unique O-glycopeptides with localized O-glycosites, and O-Pair Search identified 354 localized O-glycosites in total when allowing 5 glycans per peptide from the 12-glycan database.

In all, we show that O-Pair Search can reduce O-glycopeptide search times by >2000x over the most widely used commercial glycopeptide search tool, Byonic. Additionally, O-Pair Search identifies more O-glycopeptides than Byonic and provides O-glycosite localizations using graph theory and localization probabilities. O-Pair Search also introduces a novel classification scheme to unify data reporting across the glycoproteomic community. These Localization Levels are automatically calculated by O-Pair Search to indicate if all (Level 1), at least one (Level 2), or none (Level 3) of the O-glycosites are confidently localized. We further demonstrate the utility of O-Pair Search by searching a large published dataset of urinary O-glycopeptides, significantly increasing the number of glycopeptides identified and providing site-specific localization for >350 O-glycosites.

## METHODS

### O-Pair Search Algorithm

O-Pair Search has been implemented in MetaMorpheus^12^, an open-source search software useful for a variety of different applications including: bottom-up, top-down, PTM discovery, crosslink analysis and label free quantification. O-Pair is optimally designed for identifying O-glycopeptides from tethered collision- and electron-based dissociation spectra collected from the same precursor ion. However, it is also capable of identifying O-glycopeptides from spectra obtained using other fragmentation schemes and modalities. O-Pair Search occurs in three stages: (**Fig. 1a**) 1) identification of peptide candidates using an ion-indexed open search; 2) localization of O-glycosites with a graph-based localization algorithm; and 3) calculation of site-specific localization probabilities. Upon completion of these stages, the O-glycopeptide localization levels (**Fig. 1d**) are determined and reported along with the false discovery rates (FDR), which are presently estimated using the target-decoy strategy.

#### 1. Ion-indexed open search

MetaMorpheus uses ion-indexed open search^13^ to quickly identify peptide candidates for each spectrum. O-glycosylation is a labile modification and O-glycopeptides under collision-based dissociation in mass spectrometry generate peptide backbone fragment ions rarely retaining the glycans. Thus, even though an O-glycopeptide can be modified with multiple O-glycans, an O-glycopeptide HCD spectrum could be searched to determine the amino acid backbone without considering the O-glycans.

In an ion-indexed open search, a lookup table is created that includes a complete set of theoretical target and decoy fragment masses from the entire protein database, each labeled with the peptide from which it is derived. A collection of all peptides with fragments matching any peak in a given MS2 spectrum is assembled. The peptide candidates are then chosen from those peptides with the most matching fragments. The usage of an ion-indexed algorithm avoids unnecessary comparisons between experimental and theoretical spectra and makes it unnecessary to consider the variety of posttranslational modifications that might be present. The peptide candidates with the highest scores are retained for glycan identification and localization.

For each peptide candidate retained from the open search, the mass difference between the unmodified peptide backbone and the experimental precursor mass is computed. The mass difference is hypothesized to be the sum of all glycan masses on the peptide. We refer to the collection of glycans on a given peptides as the glycan group: *mass of glycan group = precursor mass - peptide mass*. All glycan groups whose mass equals the mass difference within the specified mass tolerance are considered as glycan group candidates for glycosite localization.

#### 2. Graph-based localization

The graph algorithm is specially optimized for O-glycosite localization. A directed acyclic graph is constructed to represent all possible O-glycan modified forms of a peptide candidate and each of its corresponding glycan group candidates. If a peptide candidate corresponds to several different glycan group candidates within the mass tolerance limitation, several graphs are constructed.

The graph is constructed from left to right, beginning with a ‘Start’ node at the N-terminal side of the peptide and ending with an ‘End’ node at the C-terminal side. Nodes, vertically aligned, are added to the graph for each corresponding serine or threonine because these amino acids are the only two allowed for O-glycosite occupancy. One vertical node designates the site as unoccupied and is labeled with ‘N’. Vertical nodes are then added, one for each potential glycan at the current position. These are labelled ‘A’, ‘B’ and so on. Additional vertical nodes are added representing combinations of glycans that may have occurred for the portion of the peptide represented by that vertical column of the graph. Combination nodes are labelled, for example. ‘A+B’. These nodes and labels are repeated at each serine and threonine. Next, adjacent nodes are connected by edges representing the accumulation of glycans across the peptide backbone. Nodes that are not possible given the constraints of the total peptide mass, which stipulate the number and kinds of glycans on the peptide remain disconnected. This process culminates in a graph representing all possible glycopeptides, where each individual continuous path from Start to End represents one unique glycopeptide.

Next, we associate theoretical fragment ions with each node. Here we need to make clear which amino acids and glycans from the peptide are included. Beginning at the N-terminus, the node represents the peptide up to AND INCLUDING the amino acid listed for the node. Beginning at the C-terminus, the node represents the peptide up to BUT NOT INCLUDING the amino acid listed for the node. The two portions of a peptide associated with a node are complementary to each other and do not cross over. Each node has associated with it all possible theoretical peptide fragment masses whose accumulated mass can be uniquely attributed to the glycopeptide segment containing the amino acids up to that point. The MetaMorpheus score for the entire peptide is the count of matching fragments from all nodes in the path plus the fraction of spectrum intensity attributable to the matched fragments. The glycopeptide with the highest MetaMorpheus score can be extracted with dynamic programming and is designated as the match and reported in the results.

We provide the hypothetical example illustrated in **Fig.1b** to aid understanding of the graph theoretical model. The example O-glycopeptide contains 8 O-glycosites. The glycan group consists of two glycans ‘A’ and ‘B’. Either of the two glycans can occupy any one of the eight positions subject to the following requirements: a maximum of two glycans can be on the peptide, only one glycan is allowed per position; and each glycan can appear only once on a given peptide. For this example, there are 56 total (**Supplementary Table 2**) different modified forms in the graph. The weight of nodes vertically aligned is determined by the number of associated theoretical fragment ions. In the example, the nodes associated with amino acid S9 can be matched to theoretical fragments c9, c10, c11, z9, z10, z11. The path highlighted in orange represents that the peptide is modified on S9 with glycan A and S12 with glycan B.

#### 3. Site-Specific Localization Probability

We use an iterative method to track the localization scores from all the potential paths of the graph to calculate site specific localization probability of a glycoPSM. These scores are integrated with a random event-based localization method similar to a method described previously in PhosphoRS^18^. The integer part of the localization is the MetaMorpheus score, k, which is the total number of matched peaks. This is applied to a cumulative binomial distribution for calculating probability P as follows:

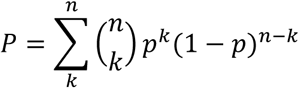

In the formula, *n* is the number of theoretical fragment ions; *p* is the probability of randomly matching a single theoretical fragment ion given specified tolerances.

One significant difference from PhosphoRS is that the extracted peak depth is not optimized to achieve maximal differentiation. Finally, localization level is assigned by considering the ambiguity of paths, the matched fragment ions corresponding to each localized O-glycosite and the site-specific probabilities.

### Data Analysis

All searches were performed on a PC running Windows 10 Education (version 1909), with two 2.20 GHz Intel Xeon Silver 4114 CPU processors with 64 GB of installed RAM. Up to 40 virtual processors were available to use for searching. Generally, 16 cores were used per search, but variations were used as described in the text. An O-glycan database of 12 common O-glycans was used for all searches^8^, except for the 32-glycan database used for the urinary O-glycopeptide dataset as described in **Fig. 2l**, which was compiled using literature sources.^32,37^ Both glycan databases are provided as supplementary data. Data from these analyses are available in the Supplementary Information. A FASTA database of the four standard mucins used in the literature data (CD43, MUC16, PSGL-1, and Gp1ba) were used for all searches unless otherwise noted, and known signaling peptide sequences were removed from the FASTA entries.

### Byonic Searching

The standalone Byonic^23^ environment (v 3.7.4, Protein Metrics) was used for all searches of the mucin O-glycopeptide dataset^11^, where the maximum allowed cores is 16. O-glycan modification from the 12 O-glycan database was set to common2, common3, or common4, as indicated in the text (meaning they could occur 2, 3, or 4 times, respectively, on the same glycopeptide). The total common max value was set to match the value used for O-glycans, and the total rare max was set to 1. Other modifications were: carbamidomethyl at cysteine (+57,021644, fixed), oxidation at methionine (+15.994915, common2), and deamidation at asparagine (+0.984016, rare1). A FASTA file of the four mucin standards was used as the protein database, with reverse sequences appended as decoys by Byonic. See **Supplementary Note 2** for more discussion about databases. Cleavage specificity was set as fully semi-specific for C-terminal to R and K residues (i.e., semi-tryptic) with two missed cleavages allowed. Precursor mass tolerance was set to 10 ppm with fragment mass tolerance(s) set to 20 ppm. Fragmentation was set to HCD & EThcD for appropriate raw files, and protein FDR was set to 1%. Byonic results were processed as described in ref 11. Briefly, following each search, peptide spectral match (PSM) lists were exported as .csv files from the Byonic viewer using all columns. Filtering Byonic search results is necessary to retain only high-quality identifications and minimize false positives^30^; here, filtering metrics included a Byonic score greater than or equal to 200, a logProb value greater than or equal to 2, and peptide length greater than 4 residues. The relaxed filtering metrics (**Supplementary Fig. 3**) used a score filter of 50 or higher and a required logProb value greater than or equal to 1.

### O-Pair Search

O-Pair Search was performed in MetaMorpheus (0.0.307), which is available at https://github.com/smith-chem-wisc/MetaMorpheus. O-Pair Search is designed to be used with high-resolution data^38^. The “Glyco Search” option was selected, where the O-glycopeptide search feature was enabled and the Oglycan.gdb glycan database was selected, representing the same 12 common O-glycan database used above. The “Keep top N candidates” feature was set to 50, and Data Type was set as HCD with Child Scan Dissociation set as EThcD. The “Maximum OGlycan Allowed” setting was varied as discussed in the text, where this number represents both the maximum number of O-glycan modifications that could occur on a glycopeptide candidate and the number of times each O-glycan could occur per peptide. For the majority of searches following the results obtained in Fig. 2g, the Maximum Oglycan Allowed” was set to 5 unless otherwise noted. Under Search Parameters, both “Use Provided Precursor” and “Deconvolute Precursors” were checked. Peak trimming was not enabled and Top N peaks and minimum ratio were set to 1000 and 0.01, respectively. In-Silico Digestion Parameters were set to generate decoy proteins using reversed sequences, and the initiator methionine feature was set to “Variable”. The maximum modification isoforms allowed was 1024, and the minimum and maximum peptide length values were set to 5 and 60 respectively. The protease was set to semi-trypsin with 2 missed cleavages allowed, unless otherwise noted (**Supplementary Fig. 4**). The number of database partitions was set to 1 unless noted below. Precursor and product mass tolerances were 10 and 20 ppm, respectively, and the minimum score allowed was 3. The maximum number of threads, i.e., cores, was varied as described in the text, with 16 cores being the default used in this study unless otherwise noted. Modifications were set as Carbamidomethyl on C as fixed, and Oxidation on M and Deamidation on N as variable.

O-Pair Search produces two separate PSM files, one for non-glycopeptides and one for glycopeptides. The numbers of non-glycopeptide identifications were calculated by filtering the single_psm file to include only target PSMs (T) with q-values less than 0.01. The same target and q-value filterings were used for O-glycopeptide identifications in the glyco_psm file. Localization Level assignments were calculated using the provided outputs following target and q-value filtering, and all were confirmed manually for data represented in **Fig. 2a-f**. The UpSet plot in Supplementary Fig. 5 was made using https://asntech.shinyapps.io/intervene/^39^.

Entrapment databases used for **Fig. 2h** and **2i** were compiled from several different sources. The canonical mucin database (20 entries) was compiled using annotated mucins available at http://www.medkem.gu.se/mucinbiology/databases^40^. The FBS database (86 entries) was generated from data provided from Shin et al^41^. The database of CD markers, i.e., cluster of differentiation markers known to be cell surface molecules, was downloaded from the Human Protein Atlas (https://www.proteinatlas.org/)^42^. The E. coli, yeast, and mouse proteome databases were retrieved from the Uniprot Consortium^43^. Sequences for the four mucin standards in the mixture that was analyzed were appended to each. See **Supplementary Note 2** for more discussion about the databases used. For searches performed with each of these databases, the Number of Database Partitions was set to 16, and 16 cores were also used for each search. The false discovery rate was calculated after filtering for target hits and q-value < 0.01 in the glyco_psms file, by taking the ratio of the total number of GlycoPSMs that did not originate from the four mucin standard proteins (false positives) to the total number of GlycoPSMs. This was performed when filtering based on Localization Levels as indicated in the text.

### Analysis of Urinary O-glycopeptide Dataset

Raw data is available for download from MassIVE (identifier MSV000083070) as provided in ref 32, and processed data for part of this dataset (Fraction 1 and Fraction 2) is available in ref 31. As described in the Supplemental Material in ref 31, raw files 170919_11.raw, 170921_06.raw, and 170922_04.raw correspond to Fraction 1. Raw files 170919_08.raw and 170921_03.raw are the only two files available for download from MassIVE that are from Fraction 2. We processed those sets of three and two files as Fraction 1 and Fraction 2, respectively, and then processed all ten files available for download from MassIVE, as indicated in **Fig. 2l**. Identifications from Protein Prospector and Byonic provided in the supplemental material from ref 31 were used from all three search conditions provided (described in detail in ref 31), with duplicate identifications between the searches removed. To convert Protein Prospector identifications to our Localization Levels scheme, all identifications containing “@” but not “|” were classified as Level 1 or 1b, because “@” indicates a modification assigned at a specific residue while “|” indicates an ambiguous assignment. Level 2 identifications were then added by included GlycoPSMs that included an “@”, whether or not other characters indicating ambiguity were present because “@” meant at least one modification was localized.

## Supporting information

SuppData_NumOfGlycans_GlycoPSMs

SuppData_NumOfGlycans_Glycosites

SuppData_NumOfGlycans_NonModPSMs

SuppData_EntrapmentDBs_GlycoPSMs

SuppData_EntrapmentDBs_Glycosites

SuppData_EntrapmentDBs_NonModPSMs

SuppData_Fragmentation_GlycoPSMs

SuppData_Fragmentation_Glycosites

SuppData_Fragmentation_NonModPSMs

SuppData_UrinaryOglyco_GlycoPSMs

SuppData_UrinaryOglyco_Glycosites

SuppData_UrinaryOglyco_NonModPSMs

## DATA AVAILABILITY

The data used in this manuscript are available through the Proteome-Xchange Consortium via the PRIDE partner repository^44^ with the dataset identifier PXD017646 (ref 11) and via MassIVE (http://massive.ucsd.edu) with identifier MSV000083070 (ref 32). Processed data using Byonic and Protein Prospector for the urinary O-glycopeptide data set was downloaded from ref 29.

## CODE AVAILABILITY

O-Pair Search is available in MetaMorpheus (0.0.307), which is open-source and freely available at https://github.com/smith-chem-wisc/MetaMorpheus under a permissive license. All source code was written in Microsoft C# with .NET CORE 3.1 using Visual Studio.

## ACKNOWLEDGMENTS

We greatly appreciate discussions with Zach Rolfs, Robert J. Millikin and other Smith group members to enhance software analysis speed and address challenges in implementing ideas. This work was supported by National Institute of Health (NIH) Grant R35 GM126914 awarded to L.M.S. and R01 GM59907 awarded to C.R.B, as well as with support from the Howard Hughes Medical Institute. N.M.R. was funded through an NIH Predoctoral to Postdoctoral Transition Award (Grant K00 CA21245403).

## AUTHOR INFORMATION

### Contributions

L.L. and N.M.R. contributed equally to this work. L.L. conceived the project and software design, wrote software, analyzed data, and wrote the paper. N.M.R. conceived the project and software design, advised on software development, analyzed a majority of data, and wrote the paper. M.R.S. designed software and supervised the project. C.R.B. and L.M.S. supervised the project. All authors discussed results and edited the paper.

## SUPPLEMENTARY INFORMATION

Supplementary Information includes Supplementary Figs. 1-12, Supplementary Tables 1 and 2, and Supplementary Notes 1 and 2:

**Supplementary Fig. 1**: Potential glycosites per theoretical peptide digested from standard mucin proteins in this study

**Supplementary Fig. 2**: Comparing Byonic and O-Pair Search when allowing 2 glycans per peptide

**Supplementary Fig. 3**: Comparing Byonic and O-Pair Search when relaxing Byonic filtering metrics for a 3-glycans-per-peptide search

**Supplementary Fig. 4**: Overlap of identifications when allowing for more glycans per peptide

**Supplementary Fig. 5**: Non-modified peptide identifications

**Supplementary Fig. 6**: Elution times correlate for related glycoforms of the same peptide sequence

**Supplementary Fig. 7**: Visualizing eluting isoforms of localized glycopeptides

**Supplementary Fig. 8**: Re-evaluating ETD and EThcD fragmentation data using O-Pair Search

**Supplementary Fig. 9**: Search speed benefits with O-Pair Search remain even with fewer cores

**Supplementary Fig. 10**: Identification of O-glycosites in four standard mucins

**Supplementary Fig. 11**: Comparing O-Pair Search with Byonic and Protein Prospector for Fraction 2 of the urinary O-glycopeptide dataset.

**Supplementary Fig. 12**: Comparing computational complexity (in Supplementary Note 1)

**Supplementary Note 1**: Computational complexity analysis

**Supplementary Note 2**: Entrapment database generation

**Supplementary Table 1**: Computation complexity analysis (in Supplementary Note 1)

**Supplementary Table 2**: Comparing computational complexity (in Supplementary Note 1)

Also included are 12 Supplementary Data files:

OPairSearch_EntrapmentDatabases_GlycoPSMs.xlsx

OPairSearch_EntrapmentDatabases_Glycosites.xlsx

OPairSearch_EntrapmentDatabases_NonModifiedPSMs.xlsx

OPairSearch_FragmentationTest_GlycoPSMs.xlsx

OPairSearch_FragmentationTest_Glycosites.xlsx

OPairSearch_FragmentationTest_NonModifiedPSMs.xlsx

OPairSearch_NumberOfGlycans_GlycoPSMs.xlsx

OPairSearch_NumberOfGlycans_Glycosites.xlsx

OPairSearch_NumberOfGlycans_NonModifiedPSMs.xlsx

OPairSearch_UrinaryOglycopeptides_GlycoPSMs.xlsx

OPairSearch_UrinaryOglycopeptides_Glycosites.xlsx

OPairSearch_UrinaryOglycopeptides_NonModifiedPSMs.xlsx

