## Supplementary Information for "O-Pair Search with MetaMorpheus for O-glycopeptide Characterization"

This Supplementary Information includes Supplementary Figs. 1-12, Supplementary Notes 1 and 2, and Supplementary Tables 1 and 2:

**Supplementary Fig. 1**: Potential glycosites per theoretical peptide digested from standard mucin proteins in this study
**Supplementary Fig. 2**: Comparing Byonic and O-Pair Search when allowing 2 glycans per peptide
**Supplementary Fig. 3**: Comparing Byonic and O-Pair Search when relaxing Byonic filtering metrics for a 3-glycans-per-peptide search
**Supplementary Fig. 4**: Overlap of identifications when allowing for more glycans per peptide
**Supplementary Fig. 5**: Non-modified peptide identifications
**Supplementary Fig. 6**: Elution times correlate for related glycoforms of the same peptide sequence
**Supplementary Fig. 7**: Visualizing eluting isoforms of localized glycopeptides
**Supplementary Fig. 8**: Re-evaluating ETD and EThcD fragmentation data using O-Pair Search
**Supplementary Fig. 9**: Search speed benefits with O-Pair Search remain even with fewer cores
**Supplementary Fig. 10**: Identification of O-glycosites in four standard mucins
**Supplementary Fig. 11**: Comparing O-Pair Search with Byonic and Protein Prospector for Fraction 2 of the urinary O-glycopeptide dataset.
**Supplementary Fig. 12**: Comparing computational complexity (in Supplementary Note 1)

**Supplementary Note 1**: Computational complexity analysis
**Supplementary Note 2**: Entrapment database generation

**Supplementary Table 1**: Computational complexity analysis (in Supplementary Note 1)
**Supplementary Table 2**: Comparing computational complexity (in Supplementary Note 1)

Also included are 12 Supplementary Data files:
OPairSearch_EntrapmentDatabases_GlycoPSMs.xlsx
OPairSearch_EntrapmentDatabases_Glycosites.xlsx
OPairSearch_EntrapmentDatabases_NonModifiedPSMs.xlsx
OPairSearch_FragmentationTest_GlycoPSMs.xlsx
OPairSearch_FragmentationTest_Glycosites.xlsx
OPairSearch_FragmentationTest_NonModifiedPSMs.xlsx
OPairSearch_NumberOfGlycans_GlycoPSMs.xlsx
OPairSearch_NumberOfGlycans_Glycosites.xlsx
OPairSearch_NumberOfGlycans_NonModifiedPSMs.xlsx
OPairSearch_UrinaryOglycopeptides_GlycoPSMs.xlsx
OPairSearch_UrinaryOglycopeptides_Glycosites.xlsx
OPairSearch_UrinaryOglycopeptides_NonModifiedPSMs.xlsx

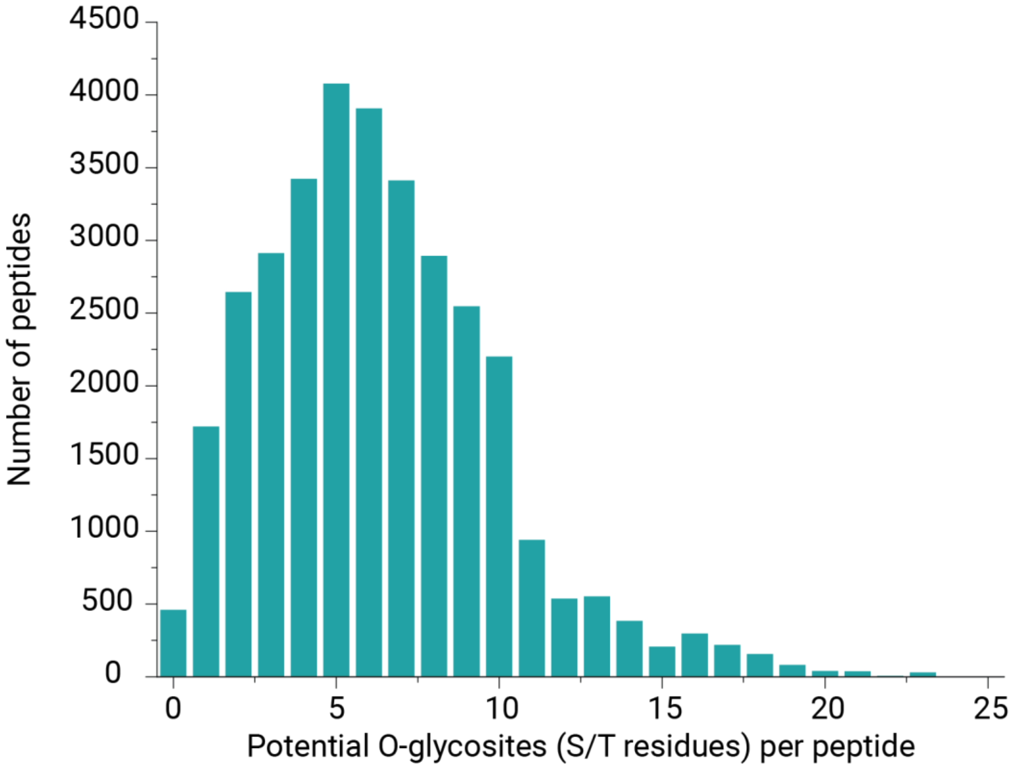


**Supplementary Figure 1**. **Potential glycosites per theoretical peptide digested from standard mucin proteins in this study.** On average, peptides derived from mucins can have approximately 6 serine/threonine residues.

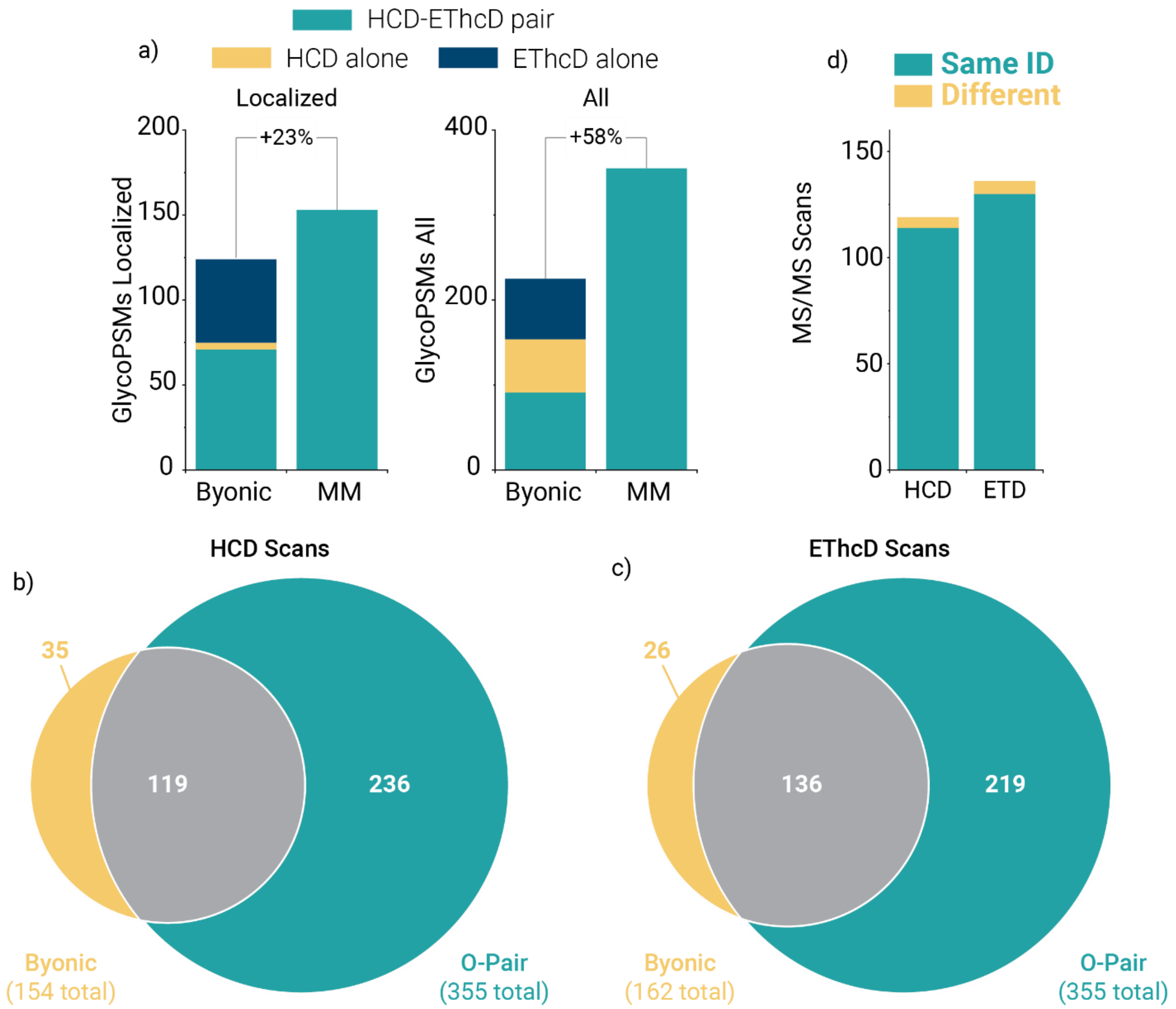


**Supplementary Figure 2. Comparing Byonic and O-Pair Search when allowing 2 glycans per peptide. a)** O-Pair searching identifies more localized and total (all) GlycoPSMs when allowing for 2 glycans per peptide from a 12-glycan database. Byonic identifications are grouped into HCD-EThcD pairs (where paired scans identified the same O-glycopeptide), HCD alone, and EThcD alone. The latter two cases are where an identification came only from an HCD scan or EThcD scan, but the other spectrum in the pair did not return a hit. O-Pair Search identified the majority of scans that Byonic returned as GlycoPSMs for both **b)** HCD and **c)** EThcD scans, and **d)** the overwhelming majority (~95%) of the shared identified scans mapped to the same glycopeptide.

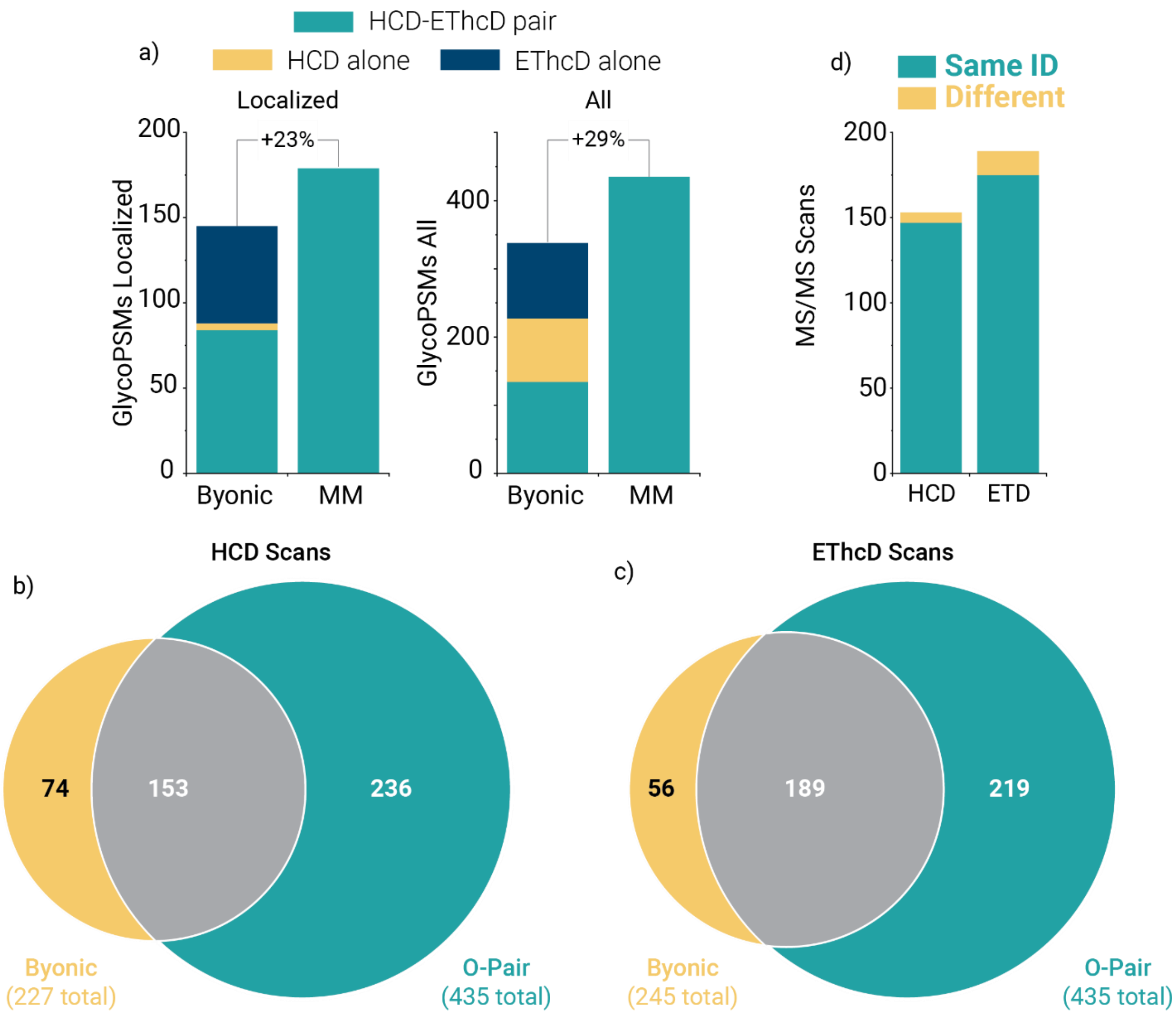


**Supplementary Figure 3. Comparing Byonic and O-Pair Search when relaxing Byonic filtering metrics for a 3-glycans-per-peptide search**. **a)** O-Pair searching identifies more localized and total (all) GlycoPSMs when allowing for 3 glycans per peptide from a 12-glycan database. Byonic identifications were filtered for scores > 50 and logProb values > 1 (compared to 200 and 2, respectively, as presented in the main text **Fig. 2**). are grouped into HCD-EThcD pairs (where paired scans identified the same O-glycopeptide), HCD alone, and EThcD alone. The latter two cases are where an identification came only from an HCD scan or EThcD scan, but the other spectrum in the pair did not return a hit. O-Pair Search identified the majority of scans that Byonic returned as GlycoPSMs for both **b)** HCD and **c)** EThcD scans, and **d)** the overwhelming majority (~95%) of the shared identified scans mapped to the same glycopeptide. The majority of additional HCD identifications from the relaxed Byonic filtering are not those that overlap with O-Pair Search identifications, while the overlapping identifications increased more than non-overlapping identifications for EThcD scans. This further supports previous findings that EThcD scans are underscored relative to HCD scans in Byonic, with HCD scans having a greater likelihood of being assigned to less confident identifications. This observation also further highlights the benefits of using HCD and EThcD spectral pairs when assigning identifications, as O-Pair Search does.

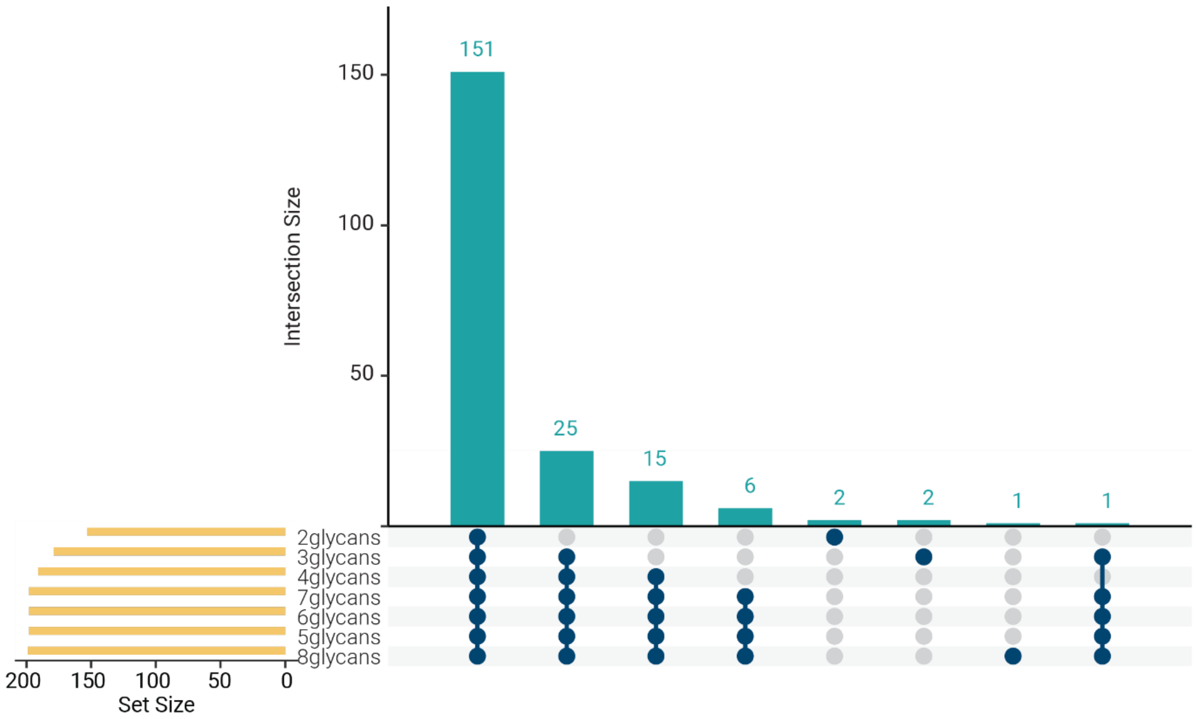


**Supplementary Figure 4. Overlap of identifications when allowing for more glycans per peptide**. The UpSet plot, which functions as a multi-dimensional Venn diagram depicting intersection size between different group as the top bar graph, shows that the core number of identifications stays the same between searches that allow for more glycans per peptide, with additional identifications remaining in the intersection of searches with progressively more glycans allowed. The total identifications count (Level1 and 1b) for each search are shown as “set size” in the bar graph in the bottom left corner.

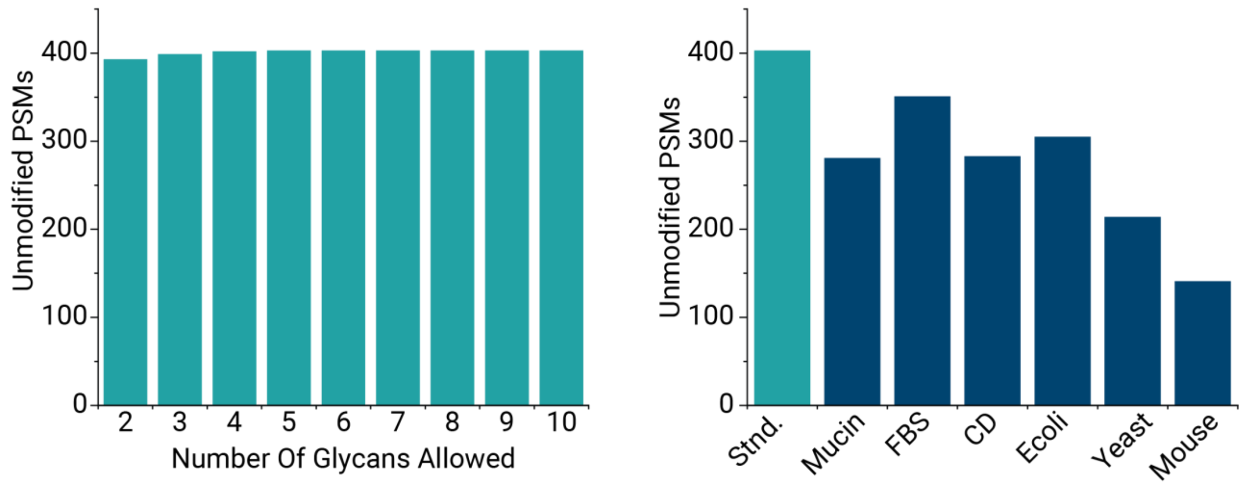


**Supplementary Figure 5. Non-modified peptide identifications**. O-Pair Search also returns identifications for non-modified peptides, i.e., standard peptide spectral matches (PSMs) that include oxidized methionine variable modifications. The number of non-modified PSMs are shown for a) the searches for the variable number of O-glycans allowed per peptide and b) the searches with six entrapment databases (with the database of 4 standard mucins provided as a reference in aqua). The size/complexity of the entrapment database caused fewer identifications for non-modified peptides, similar to O-glycopeptide identification (**Fig. 2h**).

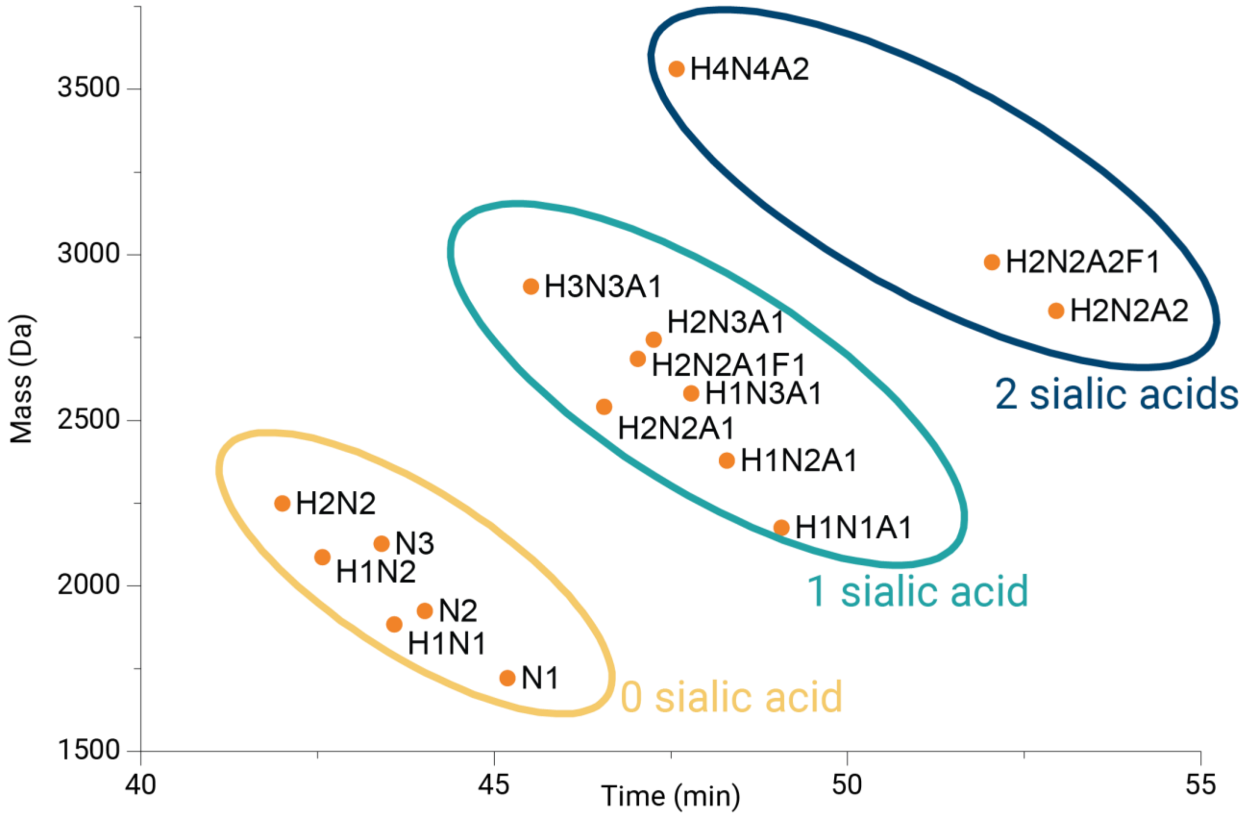


**Supplementary Figure 6. Elution times correlate for related glycoforms of the same peptide sequence**. The peptide “GLFIPFSVSSVTHK” from PSGL-1 was identified multiple times with different glycan compositions. The retention times cluster with the feature of glycans. The glycopeptides are grouped by the number of sialic acids. Glycans that contain more sialic acids elute later (between different groups); glycans that contain more neutral sugars tend to elute earlier (within one group).

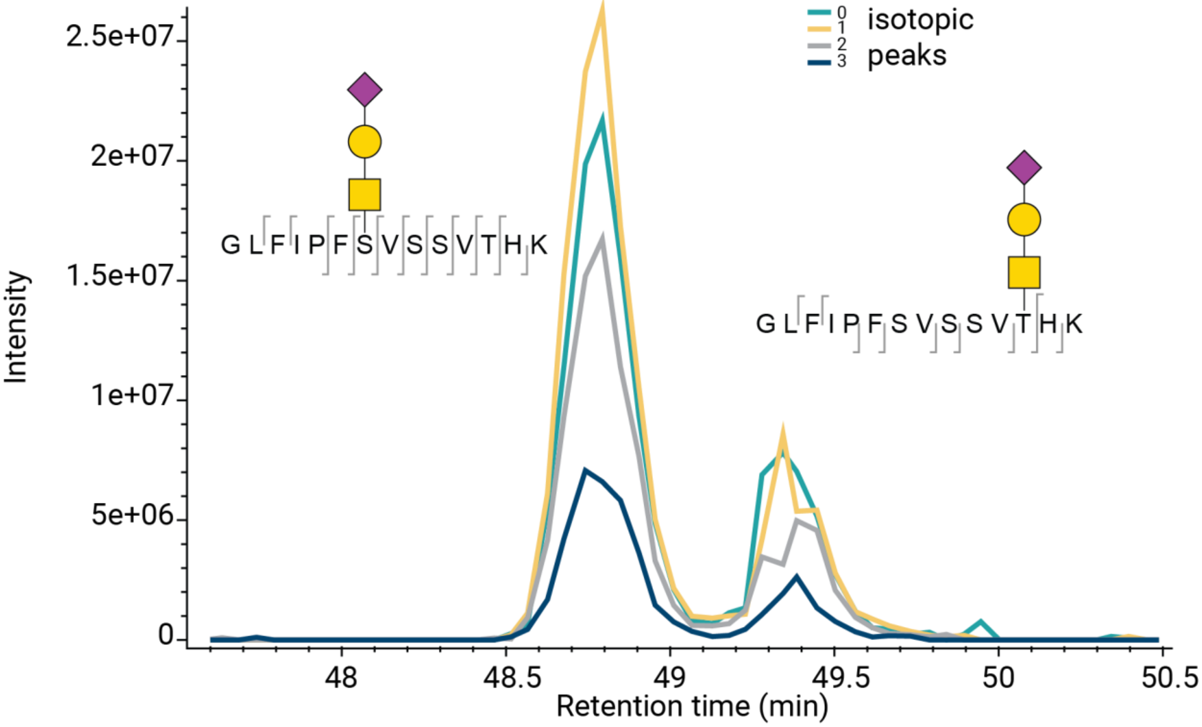


**Supplementary Figure 7. Visualizing eluting isoforms of localized glycopeptides.** Two glycopeptides, both with the aggregate mass of "GLFIPFSVSSVTHK-(H1N1A1)", are positional isoforms identified with O-Pair Search. The presence of two different elution profiles supports the presence of the two different glycoforms localized by O-Pair Search. The extracted ion current for the mass is shown for the four most abundant isotopic peaks for each glycoform.

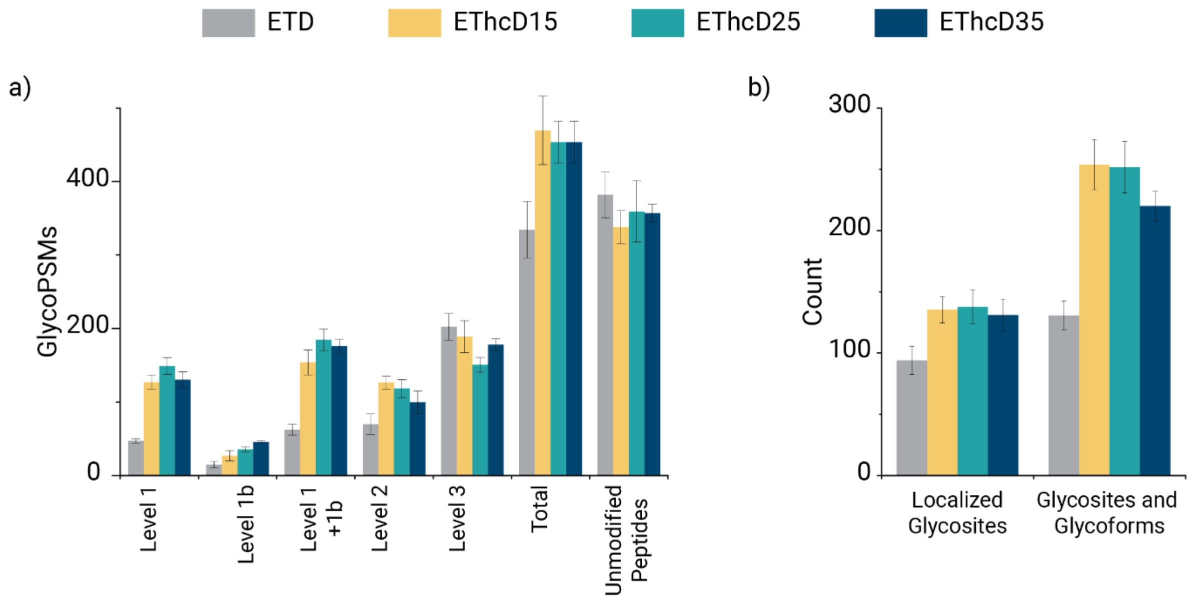


**Supplementary Figure 8**. **Re-evaluating ETD and EThcD fragmentation data using O-Pair Search.** Comparisons of electron-driven dissociation methods investigated in Main Text ref 11 are shown using O-Pair Search with Localization Level information. GlycoPSMs are shown in **a)**, including the combined Level 1 and 1b identifications, the total number of identifications, and the number of non-modified peptides detected per method. Unsurprisingly, ETD has a much higher proportion of Level 3 identifications relative to Level 1 than the EThcD methods, because non-dissociative electron transfer prevents the generation of peptide backbone fragments that are needed to localize glycosites. Even though EThcD15 generates the most total GlycoPSMs, EThcD25 has the advantage for fully localized (Level 1) identifications. **b)** The advantage in localized GlycoPSMs afforded by EThcD25 provides a slight gain in the number of localized glycosites, although the difference is minimal compared to previous analyses with Byonic. The mid-range supplemental activation energies (EThcD15 and EThcD25) provide the most localized glycosites and the most glycoforms, i.e., the combination of glycosites and the different glycans observed on them.

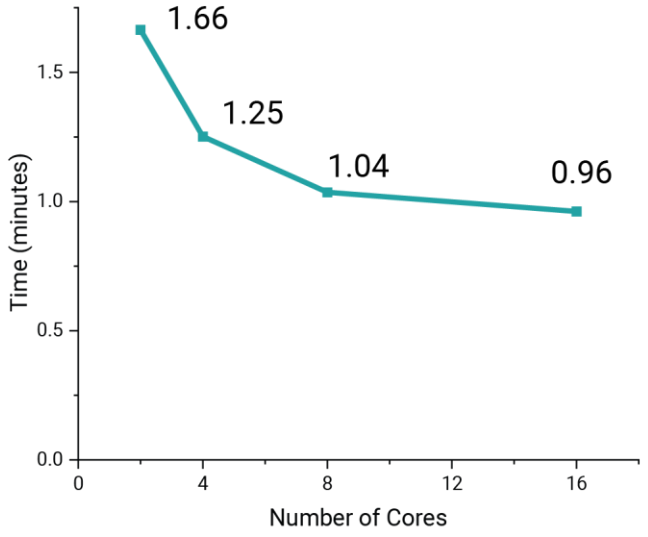


**Supplementary Figure 9. Search speed benefits with O-Pair Search remain even with fewer cores.** Here, the time needed to complete a full analysis of a raw file analogous to that shown in Fig. 2c and Fig. 2g is shown for 2, 4, 8, and 16 cores when allowing for 5 glycans per peptide with a glycan database of 12 glycans. O-Pair Search speed translates to any computer used for searching, not just hyperthreaded computers.

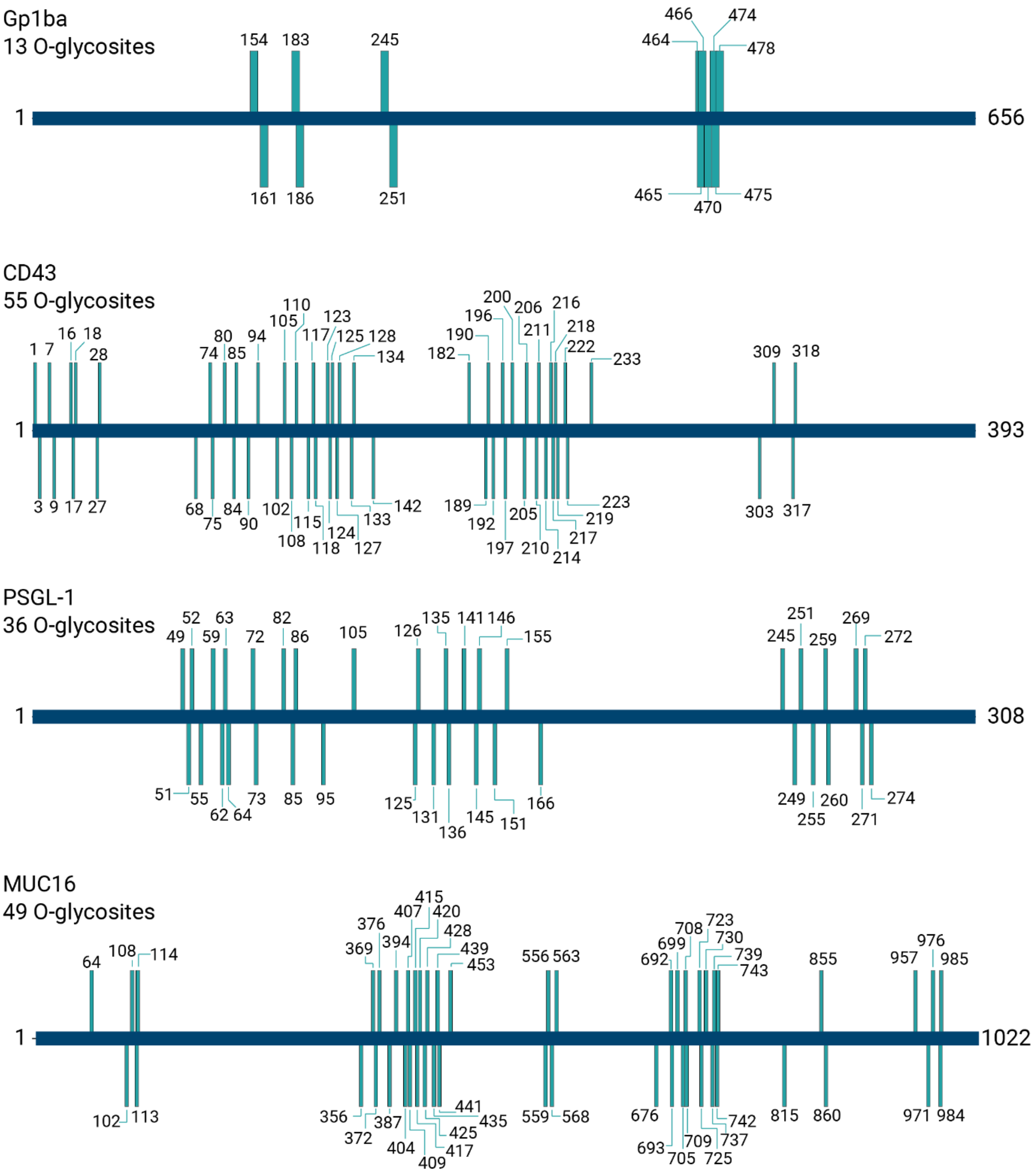


**Supplementary Figure 10**. **Identification of O-glycosites in the four standard mucins**. Searching with 5 glycans allowed from a 12 O-glycan database identifies 153 O-glycosites, with each protein extensively modified. Here, blue horizontal bars represent the protein sequences with N- and C-terminal residues numerically labeled, and green vertical bars indicate O-glycosites (residue numbers provided) localized with O-Pair Search. For comparison (but not depicted in the figure), a Byonic search with 3 glycans allowed identified 96 total O-glycosites with evidence for 41 of them as localized: 4 localized of 7 identified O-glycosites in Gp1ba, 19 localized of 37 identified O-glycosites in CD43, 12 localized of 17 identified O-glycosites in PSGL-1, and 6 localized of 35 identified O-glycosites in MUC16, Note, signal peptides were removed from the protein sequences prior to searching.

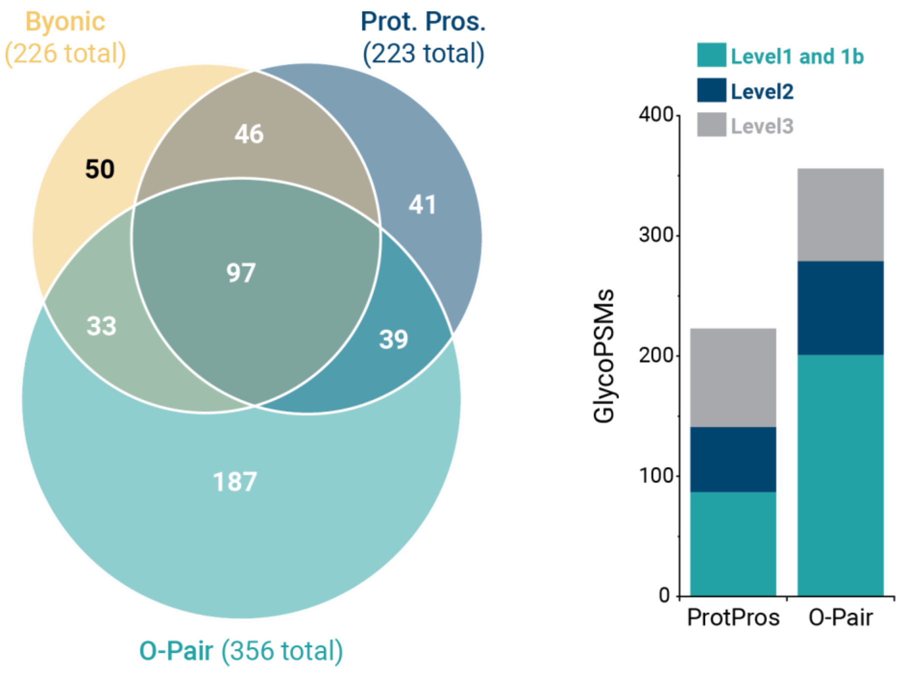


**Supplementary Figure 11**. **Comparing O-Pair Search with Byonic and Protein Prospector for Fraction 2 of the urinary O-glycopeptide dataset.** O-Pair Search has a high degree of overlap in identified spectra with the other two search algorithms while returning a greater number of unique identifications (left). We converted Protein Prospector to our Localization Level classification and compared to O-Pair Search (right).

**SUPPLEMENTARY NOTE 1**

<table>
<tr><td>Computational complexity of searching one MS2 experimental spectrum</td><td>localization</td><td>non localization</td></tr>
<tr><td>Traditional narrow search</td><td>$O\left( n * \sum_{i=0}^{m} \frac{(s)!}{(s-i)!*i!} * g^{i} \right)$</td><td>$O\left( n * \sum_{i=0}^{m} \frac{((g-1)+i)!}{(g-1)!*i!} \right)$</td></tr>
<tr><td rowspan="2">ion-indexed open search</td><td>brute force localization<br>$O\left( x + k * \frac{s!}{(s-m)!} \right)$</td><td rowspan="2">$O\left( x + k \right)$</td></tr>
<tr><td>graph-based localization<br>$O\left( x + k * d \right)$</td></tr>
</table>

**Supplementary Table 1. Computation complexity analysis**

*n* is the average number of theoretical peptides within the specified precursor mass tolerance.

*m* is the maximum number of glycosites allowed on a single peptide.

*s* is the number of S/T amino acids within a single peptide.

*g* is the number of glycan types.

*k* is the count of top peptide candidates. In general, *k* is always much smaller than *n*.

*x* is the number of peaks in an MS2 experimental spectrum, which is used to represent the constant ion indexing time and can be ignored in general.

*d* is the depth of the graph. $d = \sum_{i=0}^{m} \frac{(m)!}{(m-i)!*i!} = 2^{m}$ .

### Computational complexity analysis

We observed a great reduction in search time for O-Pair Search compared against the commercial software program Byonic for similarly configured O-glycopeptide searches. This prompted us to perform an analysis of the computational complexity of the O-Pair Search algorithm and other related methods to determine the source of this time savings.

To simplify the analysis, we treat each theoretical-experimental spectrum comparison as one-unit of computation to calculate the associated computational complexity.

#### 1) Traditional database search (narrow)

The traditional database search for O-glycopeptide identification requires the construction of a complete O-glycopeptide database prior to the matching process. The O-glycopeptide database is built using O-glycans as variable modifications. In the variable modification scenario, each peptide yields a combinatorial number of glycopeptides based the number of modifiable sites within the peptide and the number of different glycans that can occupy those sites[1]. The computational complexity (**Supplementary Table 1**) of traditional glycopeptide search is:

$$O\left( n * \sum_{i=0}^{m} \binom{s}{i} g^{i} \right)$$

#### 2) Ion-Indexed Open Search with localization

The O-Pair Search algorithm, described in detail in the Methods Section, consists of an ion-indexed open search to ascertain the amino acid sequence of the peptide backbone and a graph-theoretical approach to determining the positions of glycans adorning the peptide. We analyze below the theoretical complexity of each stage separately.

#### 2.1) Ion Indexing

Ion-indexed open search strategies have been used for bottom-up peptide identification and post-translational modification discovery[2]. Ion-indexed open searches are particularly advantageous for the latter because there is no need to select theoretical peptides based on the experimental precursor mass, which would have required construction of all theoretically modified forms. Ion indexes are generally constructed from theoretical fragments of the unmodified peptide backbone for each peptide in the database. In the case of O-pair Search, the use of an ion-indexed open search for the first stage means that there is no need to consider all the possible different glycopeptide forms that could be constructed for each peptide. A theoretical peptide database containing all possible glycans would be massive. Peptide candidates only are identified in the ion-indexed open search during the first stage. Possible glycans are considered in the next. The ion indexing process in MetaMorpheus has been heavily optimized for high speed. The computational complexity (**Supplementary Table 1**) for ion indexing is related to the number of peaks in the MS2 spectrum, which can be considered as a small constant:

$$O(x)$$

#### 2.2) Brute force localization

MetaMorpheus O-pair Search does not perform a brute force localization. However, we provide here an analysis of the brute force localization approach as a point of reference for the reader to understand it in relation to the computational complexity of the graph-based localization described immediately after. Glycan group candidates are determined after obtaining peptide candidates from the ion-indexed open search. The brute force approach to localization consists of matching each glycan group to each different peptide candidate (combinatorial), including all possible localization isomers. This process yields $\frac{s!}{(s-m)!}$ glycopeptides for consideration. The overall computational complexity (**Supplementary Table 1**) for ion-indexed open search with brute force localization is:

O ( $x + k * \frac{s!}{(s-m)!}$)

#### 2.3) Graph-based Localization

The graph-based localization was described in detail in the methods section. Here we provide additional details to compute the computational complexity. There are two sets of theoretical fragment ions associated with glycopeptides in a graph. One set of theoretical fragments are shared between multiple glycopeptides and the other set of fragments is unique to a single glycopeptide. The shared fragments are predominant. The graph-based localization saves time in constructing all possible glycopeptide forms and simultaneously enables the matching of fragments without redundancy, taking advantage of the knowledge of shared fragments.

The computational complexity is proportional to the depth $d = 2^m$ of the graph, which is a function of the number of possible glycan modifications *m*. For O-glycopeptides in general, *m* is not a large number. The overall computational complexity (**Supplementary Table 1**) for ion-indexed open search with graph-based localization is:

O ( $x + k * d$)

#### 3) Comparison

To compare the computational complexity, we ignore the values *n*, *x* and k and focus on analyzing how the numbers of glycan types and glycosites influence the computational complexity.

For example, using g = 12 glycans, s = 8 glycosites and m = 2 glycans (as in **Fig 1b**), the computational complexity (**Supplementary Table 1**) is as follows:

1) in traditional search $\sum_{i=0}^{m} \frac{(s)!}{(s-i)!*i!} * g^i = 4291$
2) in ion-indexed open search with brute force localization $\frac{s!}{(s-m)!} = 56$
3) in ion-indexed open search with graph-based localization $m^2 = 4$

Next, we look at how these numbers evolve as a function of m. We set *g* = 12 and *s* = 8 and find that the computational complexity changes dramatically for traditional narrow search and brute force methods (**Supplementary Figure 14, Supplementary Table 2**).

Using the ion-indexed open search and graph-based localization, O-Pair Search avoids the 'combinatorial explosion' that happens in for traditional searches of O-glycopeptide characterization. O-Pair Search in this way could use a large sized glycan database and expands the potential for O-glycopeptide characterization.

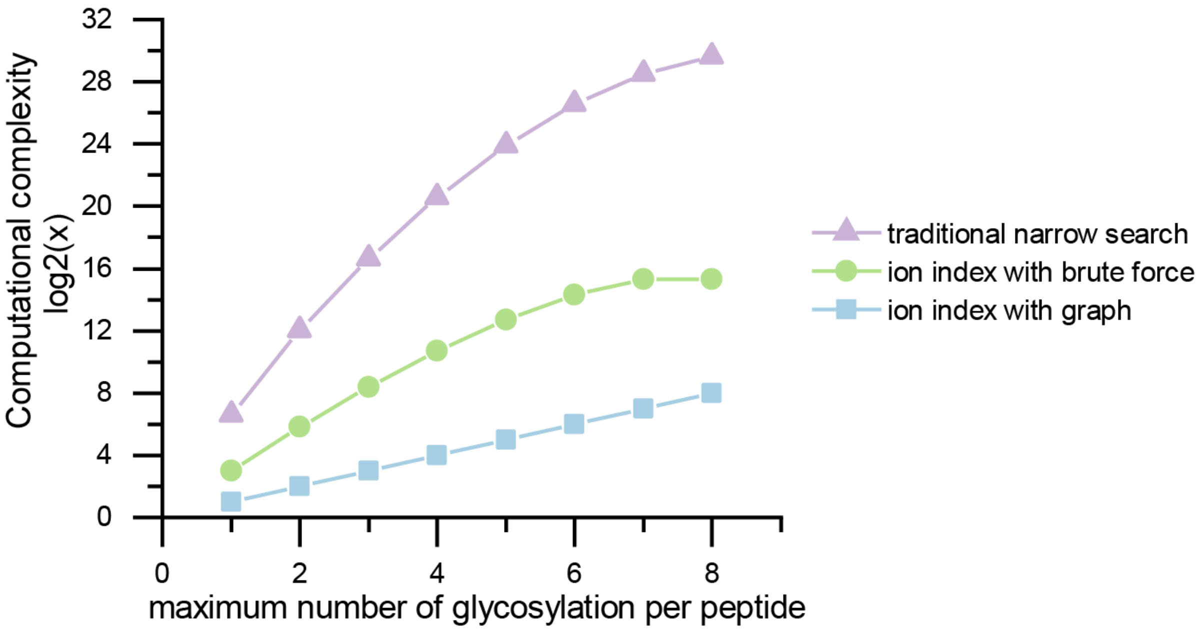


**Supplementary Figure 12.** Comparing the computational complexity in traditional narrow search, ion-indexed open search with brute force localization and graph-based localization. Note, the y-axis is on a log2 scale.

| *m* | graph | brute force | traditional |
|---|---|---|---|
| 1 | 2 | 8 | 97 |
| 2 | 4 | 56 | 4129 |
| 3 | 8 | 336 | 100897 |
| 4 | 16 | 1680 | 1552417 |
| 5 | 32 | 6720 | 1.55E+07 |
| 6 | 64 | 20160 | 9.91E+07 |
| 7 | 128 | 40320 | 3.86E+08 |
| 8 | 256 | 40320 | 8.16E+08 |

**Supplementary Table 2.** Comparing the computational complexity in traditional narrow search, ion-indexed open search with brute force localization and graph-based localization. The number of glycan types is set to $g = 12$, the number of glycosites is set to $s = 8$, and m is the maximum number of glycosylation allowed on a single peptide.

### SUPPLEMENTARY NOTE 2

Several entrapment databases were constructed to evaluate O-Pair Search performance, both for the time it takes to complete searches with progressively larger databases and for false discovery rate calculations. The goal was to create background proteomes that not only varied in size and complexity, but also established different degrees of relevance to the four mucin standard proteins that were actually present in the sample: CD43, PSGL-1, MUC16, and Gp1ba.

First, we appended the four true positive mucins to 20 canonical human mucin sequences[3]. Here, we create a “high mucin” background, where the false positives are very similar to the target proteins in sequence composition and proclivity for O-glycosylation. This also represents a close approximation of real-world scenarios where digested mucin O-glycopeptides would be screened from pools of several mucins with the need to accurately identify which glycopeptides match to which proteins. This entrapment database also represents an algorithmic challenge, where there are a multitude of O-glycopeptide candidates generated with a high number of O-glycosites.

The second entrapment database comprised the four true mucin standard sequences combined with common fetal bovine serum (FBS) proteins[4], many of which are glycosylated. This database represents a “high glycoprotein, mammalian but not human” background, where the majority of glycoproteins in the database have the capacity to be glycosylated in actuality but there should be virtually no peptides mapping to bovine sequences considering the human mucin standard protein source. This also represents a situation many researchers may encounter where FBS proteins are present due to preparation conditions and may need to be accounted for. Computationally, this is moderately more entries than the mucin background above, but with fewer overall mucin-type proteins to produce large quantities of high S/T density glycopeptide candidates.

Human cluster of differentiation (CD) markers, which are known cell surface proteins, were downloaded from the Human Protein Atlas[5] and used for the third entrapment database. This set of proteins represents a “high glycoprotein, human” background, as the majority of CD markers are known to be glycosylated. Several CD markers are classified as mucins, so this database challenges the accuracy of assignment using proteins that could potentially have real O-glycopeptide assignments while also increasing the number of entries ~5-10 fold over the first two databases.

In principle, proteins from these first three entrapment databases could have the type of O-glycosylation also seen on the true positive mucin standards. Choosing entrapment databases of the entire proteomes of E. coli and S. cerevisiae removes this constraint, as neither species should contain the glycans in the O-glycan database or peptides sequences that map to mammalian proteins. They represent “non-glycoprotein, non-species related” backgrounds that should have little in common with the true positive mucin standard proteins. They also represent significant increases in database size to ~900 and ~6700 protein entries, respectively. Thus, these two databases serve to challenge the accuracy of O-Pair Search and the search speed when considering proteome-scale data.

Finally, the entire mouse proteome represents a large entrapment database that can test both the speed of the O-Pair Search when search a database approximately analogous to the size of the human proteome while also permitting evaluation of the accuracy of assignments, as O-glycopeptides from the human mucin standards should not map to murine proteins.

Together, these six entrapment databases test the speed and accuracy of O-Pair Search, both when the background looks similar to the known mucin standards and when it looks entirely different to the mucins of interest.
